# Identification and design of vinyl sulfone inhibitors against Cryptopain-1 – a cysteine protease from cryptosporidiosis-causing *Cryptosporidium parvum*

**DOI:** 10.1101/332965

**Authors:** Arpita Banerjee

## Abstract

Cryptosporidiosis, a disease marked by diarrhea in adults and stunted growth in children, is associated with the unicellular protozoan pathogen *Cryptosporidium;* often the species *parvum*. Cryptopain-1, a cysteine protease characterized in the genome of *Cryptosporidium parvum*, had been earlier shown to be inhibited by a vinyl sulfone compound called K11777 (or K-777). Cysteine proteases have long been established as valid drug targets, which can be covalently and selectively inhibited by vinyl sulfones. This computational study was initiated to identify purchasable vinyl sulfone compounds, which could possibly inhibit cryptopain-1 with higher efficacy than K11777. Docking simulations screened a number of such possibly better inhibitors. The work was furthered to probe the enzymatic pocket of cryptopain-1, through *in-silico* mutations, to derive a map of receptor-ligand interactions in the docked complexes. The idea was to provide crucial clues to aid the design of inhibitors, which would be able to bind the protease well by making favorable interactions with important residues of the enzyme. The analyses dictated placement of ligands towards the front of the enzymatic cleft, and disfavored interactions deep within. The S1’ and S2 subsites of the enzyme preferred to remain occupied by polar ligand subgroups. Reasonably distanced ring systems and polar backbones of ligands were desired across the cleft. Large as well as inflexible subgroups were not tolerated. Double ringed systems such as substituted napthalene, especially in S1, were exceptions though. The S2 subsite, which is typically a specificity determinant in papain (C1) family cysteine proteases such as cathepsin L-like cryptopain-1, can possibly accommodate polar and hydrophobic ligand subgroups alike.

## INTRODUCTION

Cryptosporidiosis is an intestinal disease that is clinically manifested by diarrhea in adults ^[1]^ and stunted growth in children ^[2]^. The infection can persist indefinitely in immunocompromised individuals such as HIV patients, and could be fatal in the form of life-threatening diarrhea ^[3]^.

The disease is caused by unicellular protozoan parasite *Cryptosporidium*, which infects humans and animals ^[4]^ through consumption of contaminated water and/or ingestion of contaminated food products ^[5]^. The majority of infections are caused by *Cryptosporidium* species *hominis* and *parvum* ^[6] [7]^.

A cysteine protease named Cryptopain-1, characterized in the genome of *Cryptosporidium parvum* ^[8]^, most likely facilitates host cell invasion and nutritional uptake (through proteolytic degradation) ^[9] [10] [11]^. The pathogenic enzyme, being cathepsin L –like, belongs to papain-like or clan CA (family C1) cysteine protease enzymes - which in general have been of particular use as therapeutic targets against parasitic infections ^[12]^. The catalytic triad of such enzymes is constituted by Cys, His and Asn residues ^[12], [13]^. Orthologous proteases to Cryptopain-1 have been validated as drug targets *viz:* cruzain (from Chagas’ disease agent *Trypanosoma Cruzi*), rhodesain (from sleeping sickness causing *Trypanosoma brucei*), falcipain-3 (from malarial parasite *Plasmodium falciparum*), SmCB1 (from intestinal schistosomiasis causing *Schistosoma mansoni*) ^[14] [15]^ etc.

Vinyl sulfone compounds have been particularly effective inhibitors of such parasitic cysteine proteases ^[13] [14] [15] [16] [17]^. These inhibitors form a covalent bond with the active site Cys thiol to bind the proteases, thereby irreversibly blocking the enzymatic pocket. Such inhibition interferes with the pathogenic activity of the proteases that would otherwise participate in general acid-base reaction for hydrolysis of host-protein peptide bonds ^[13]^. Molecular modeling studies had previously shown that unlike serine proteases (which also cleave peptide bonds and have Ser in their active site), the catalytic His in cysteine proteases remains protonated to act as a general acid ^[18]^. Hydrogen bonding between the protonated His and the sulfone oxygen of a vinyl sulfone compound polarizes the vinyl group of the ligand to impart a positive charge on its beta carbon that eventually promotes nucleophilic attack by negatively charged Cys thiolate of the protease’s active site. Vinyl sulfone class of inhibitors are preferred over other covalent inhibitors because of its selectivity for cysteine proteases over serine proteases, relative inertness in the absence of target protease ^[18] [19]^, and safe pharmacokinetic profile ^[20]^ [21].

The peptidyl vinyl sulfones that have been co-crystallized with cysteine proteases so far reveal that the –CO-NH-backbones of the pharmacologically active compounds fit snugly in the enzymatic cleft, with the ligand sidechains (or subgroups) protruding into the different subsites of the proteases. The subgroup near the vinyl carbon that undergoes nucleophilic attack is equivalent to P1 in the inhibitor/substrate ^[13]^. Therefore, ligand sidegroups starting from the vinyl side are designated as P1, P2… that interact with the S1, S2… protease subsites. The ligand subgroups beyond the sulfonyl are referred to as P1’, P2’… and they occupy the S1’, S2’… subsites on the prime side of the enzyme (**Figure 1**). Typically, the P2-S2 interaction is the key specificity determinant in papain (C1) family cysteine proteases ^[12] [13^ like cryptopain-1.

**Figure 1:**
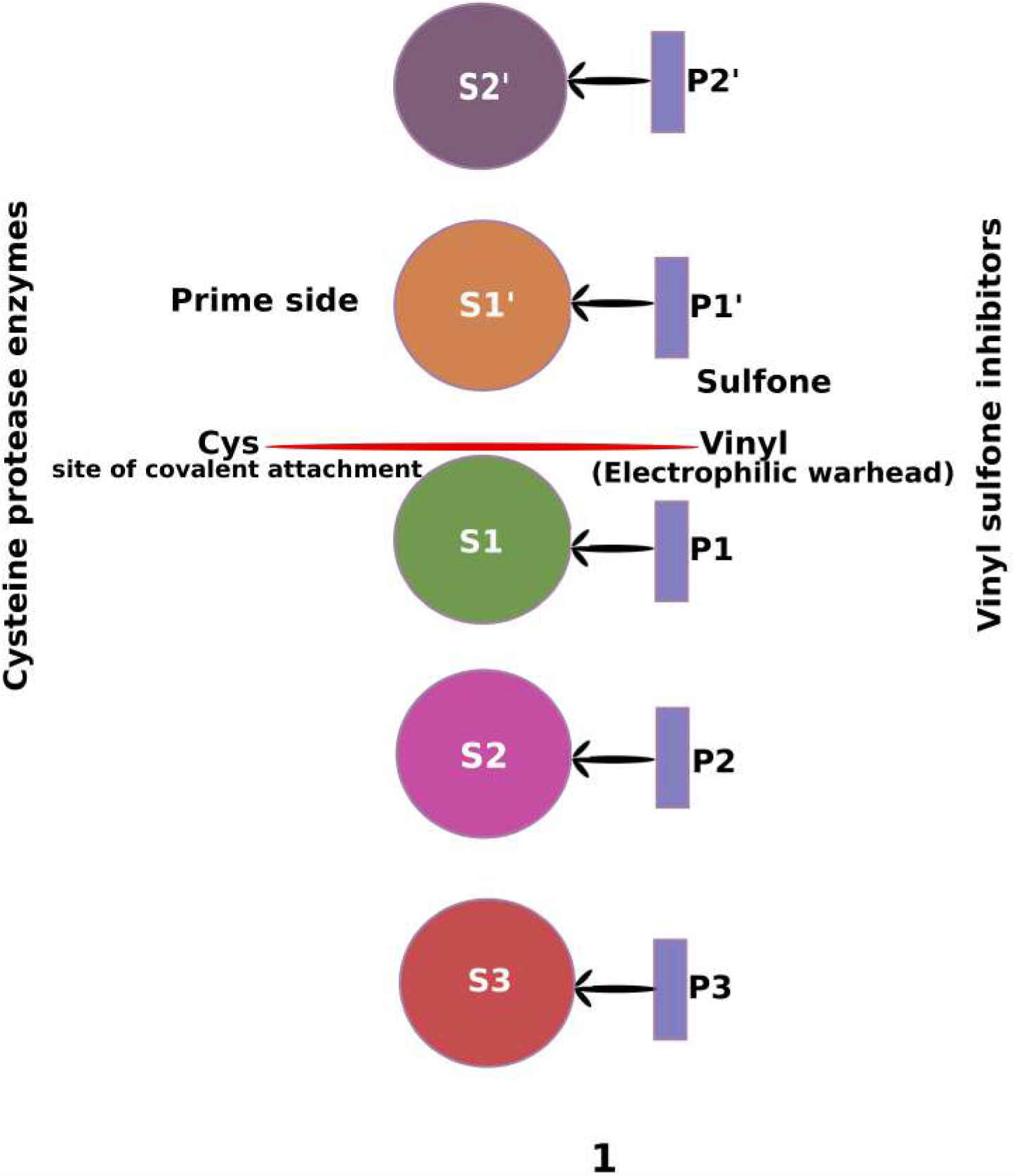
Illustration of the typical binding of vinyl sulfone inhibitors to cysteine protease enzymes. Colored spheres represent the different subsites of the enzyme, and the ligand sidechain/subgroups of the vinyl sulfone inhibitor are in violet rectangles. Spatial distribution of the subsites in three-dimensional protease structures differs from the linear arrangement that has been shown here for simplicity. The backbones of the enzyme and inhibitor are not shown. The site of covalent bond formation at C24 has been marked in red. The positioning/denotation of the ligand subgroups within the different subsites of the enzyme is according to their placement near the vinyl warhead – depicting what has been observed so far in the solved structures of peptidyl vinyl sulfone-bound cysteine proteases. The ligand sidegroup nearest the beta carbon of vinyl is P1 that fits into S1. The following ligand subgroups are P2, P3 etc. The groups beyond the sulfonyl are P1’, P2’ etc. which interact with the prime side subsites of the enzyme.

K11777 (or K-777), a vinyl sulfone that binds cryptopain-1 as its target as per inhibitor competition experiments with active site probe of the recombinant protease, has been demonstrated to arrest *Cryptosporidium parvum* growth in human cell lines at physiologically achievable concentrations ^[21]^. The cryptopain-1 structure however, by itself or in complex with K11777, has not been solved till date.

K11777-bound co-crystals of other orthologous cysteine proteases such as cruzain, rhodesain and SmCB1 ^[14] [15]^, showed the orientation of the inhibitor in the cysteine proteases as depicted in **Figure 1**. The earlier mentioned study on cryptopain-1 had simulated the binding of K11777 within the active site of the enzyme homology model ^[21]^, and mimicking nature, the inhibitor was put in an orientation as illustrated in **Figure 1**

The present computational study was initiated to explore other (purchasable) vinyl sulfones that could better bind the active site of the cryptopain-1 enzyme, with possibly higher efficacy than K11777. The study was extended to probe the enzymatic pocket of cryptopain-1 to figure preferential binding of certain ligand chemical groups at the subsites, for the purpose of providing clue to drug design against the pathogenic cysteine protease.

## MATERIALS AND METHODS

### Homology model building of enzyme

The sequence of cryptopain-1, with the accession number ABA40395.1, belonging to *cryptosporidium parvum* was retrieved from Genbank ^[22]^. The protein sequence was downloaded in fasta format.

The homology model template search for cryptopain-1 (cathepsin L-like) through NCBI BLAST against PDB database ^[23]^ led to 3F75, which is the activated *Toxoplasma gondii* cathepsin L (TgCPL) in complex with its propeptide. The template shared 48% sequence identity with the sequence to be modeled.

The homology model of cryptopain-1 was built within the full refinement module of ICM ^[24]^. The structure-guided sequence alignment between the template and the model was generated using the default matrix with gap opening penalty of 2.40 and gap extension penalty of 0.15. Loops were sampled for the alignment gaps where the template did not have co-ordinates for the model. The loop refinement parameters were used according to default settings. Acceptance ratio for the simulation process was 1.25. The generated homology model of a length of 231 amino acids was then validated in PROCHECK ^[25]^ and PROSA ^[26]^ webservers.

### Ligand structures from chemical compound database

K11777 (or K-777) was downloaded from PubChem ^[27]^ in SDF format. The vinyl sulfone substructure of K11777 was then searched in PubChem, with the additional option of ‘Ring systems not embedded’ so as to filter out those structures where the vinyl bonds would extend into ring systems. The search, which was obviously not restricted to peptidyl vinyl sulfones, led to 10,663 hits (as of April 5, 2016). 2115 compounds, which were purchasable amongst the hits, were downloaded in SDF format. The downloaded compounds were checked for redundancy. From the 1890 non-redundant vinyl sulfone compounds, 774 cyanide compounds were discarded due to the usual high toxicity profile of such compounds, and the remaining 1116 were saved to be used as ligands for docking into cryptopain-1.

### Docking simulation of covalent inhibition of enzyme

The N-terminal propeptide (which is not part of the active enzyme and acts as a self-inhibitory peptide for regulatory purposes) of the cryptopain-1 homology model was deleted. The residues were then renumbered in the enzyme model, with position 1 allocated to the beginning of the mature protease. The pdb file of the edited cryptopain-1 model was then prepared as a receptor in ICM with the addition of protons, optimization of His, Pro, Asn, Gln and Cys residues. The protonation step was crucial for mimicking the reaction (and hence bonds) between a vinyl sulfone and the cysteine protease. The active site residues of the binding pocket had been derived from the structural alignment of cryptopain-1 homology model with the orthologous cruzain that was bound to K11777 (PDB ID: 2OZ2), followed by mapping of the residues around K11777 in the cruzain onto the cryptopain-1 sequence. The pre-determined pocket residues were selected (except the catalytic Cys24 or C24) on the prepared cryptopain-1 in the GUI of ICM and the relevant box size was created on the receptor for defining the area for ligand docking. Further, C24 was selected for specifying the covalent docking site. From the set of preloaded reactions in ICM, alpha, beta-unsaturated sulfone/sulfonamide/cysteine reaction was selected, which specified the simulation of covalent bond formation between the supposedly thiolate (C24 of protease) and the beta carbon atom (of the vinyl group of ligand). The receptor maps were finally made for grid generation.

K11777, downloaded from PubChem in SDF format, was read in as a chemical table in the GUI of ICM, and was specified for docking into the prepared cryptopain-1 receptor. Thoroughness of 3.00 was set in the docking protocol, and twenty conformations of the ligand in the receptor were generated.

Following K11777, a total of 1116 non-cyanide vinyl sulfone compounds were attempted for covalent docking into the cryptopain-1 homology model, using the same protocol as described above.

### In-silico mutation of enzyme residues for assessing binding

For the purpose of evaluating the contribution of the individual residues to the binding of the ligands, mutational analysis was undertaken. The protein-ligand stability was measured by *in-silico* mutation of the contact residues in the complexes. K11777-docked cryptopain-1 and the best-scored complexes (with a score of −29 or lower) were read in separately, and then for each of them, the ligand-subgroup contacting residues were selected one at a time in the workspace panel, and were mutated to Alanine. The outputs of the calculations were displayed in several columns. dGwt column had the dG (Gibbs free energy) value for the wild type complex (without mutation), the dGmut held the dG value for the mutated complex (where the residue was mutated to Ala), and the ddGbind (dGmut – dGwt) column, which showed the binding free energy change (in Kcal/mol) upon mutation, essentially predicted the stability of the native complex, thereby hinting at the contribution of the residue in question towards binding the ligand. Positive values of ddGbind implied the mutation to be less favorable, indicating greater contribution of the wild type residue towards binding. Hence, with more positive ddGbind, better binding of the ligand by the residue could be expected. Negative values, on the other hand, implied the mutated form to be more stable, thereby delineating the native residue’s involvement in unfavorable interactions with the ligand.

The residues that were detected to make high number of favorable ligand interactions in thirty-two of the complexes (K11777-cryptopain-1 plus thirty-one best-scored ones) were subjected to a fresh round of mutations in the updated version of the ICM software. The recalculated ddGbind values were then tallied with the placement and orientation of ligand-subgroups around the residues to decipher the preference of chemical groups across the enzymatic cleft of cryptopain-1.

[The GUI of ICM was used to make the enzyme/complex structure figures. Illustration and compilation of figures were done in Inkscape, which is an open-source vector graphics editor]

## RESULTS AND DISCUSSION

### Validation of theoretical enzyme structure

The ramachandran plot for the cryptopain-1 homology model showed 98% of the residues to lie in the allowed region, and the remaining 2% to be within the generously allowed region of the plot (**Supplementary Figure 1A**). The PROSA Z-score for the cryptopain-1 model was −7.79, better than the −6.66 Z-score of its crystal structure template (**Supplementary Figure 1B**).

### Screening of docked compounds

Besides K11777, a total of 1116 purchasable, non-redundant and non-cyanide vinyl sulfone compounds were docked and scored in the cryptopain-1 homology model (50 symmetric molecules could not be docked using ICM).

Post docking, the conformation of K11777 − where the ligand P1’ group (beyond the sulfonyl) got oriented across the enzyme S1’ and its P1..P3 groups (beyond the vinyl) were placed across the S1..S3 subsites (as in **Figure 1**), and had the lowest score in the said category, was chosen as a reference for the analysis. Such orientation appeared first in the eighteenth pose (conformation) of K11777 docked into cryptopain-1, with a score of −19.15.

The conformations of some other docked vinyl sulfone compounds that had similar orientation (described above) where the ligand subgroups beyond the sulfonyl were placed across S1’ or beyond, with lowest scores <= −29.0 (and hence possibly better binders than K11777), were included in the study for further detailed analysis.

[The chemical structures of K11777 and the thirty-one best-scored vinyl sulfones are provided in **Supplementary Figure 2**, as PubChem IDs associated with (some) chemical compounds change due to frequent updates to the database. The IDs mentioned throughout the text, tables and figures are from the current PubChem records as of May 26, 2018]

**Figure 2:**
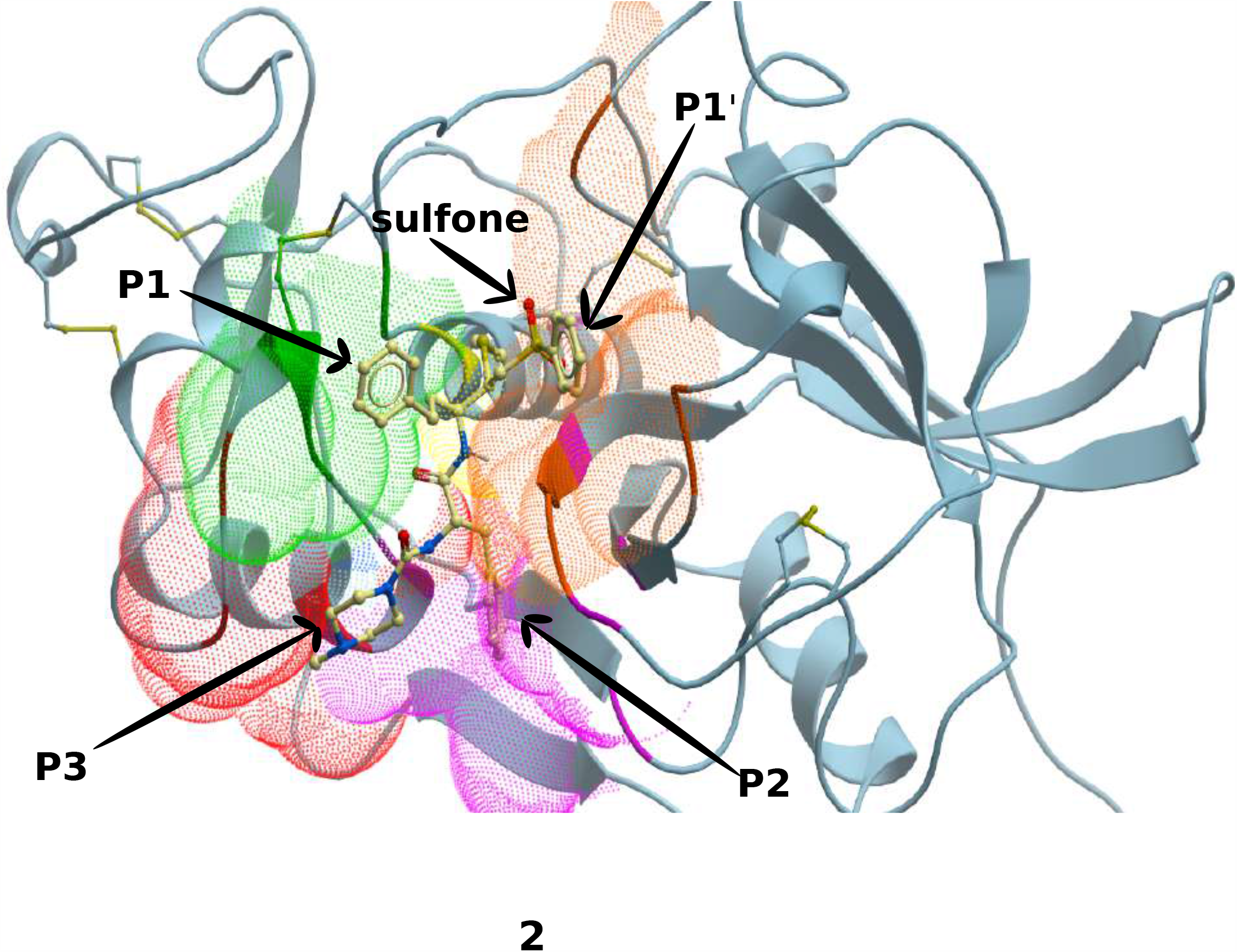
K11777 or K-777 (PubChem ID: 9851116) docked into the three-dimensional (homology) model of cryptopain-1. The selected conformation (score: −19.15) shown here conforms to the arrangement of the ligand subgoups (P1’, P1, P2, P3) in the different enzyme subsites as depicted in Figure1, and so does the color code that demarcates the subsites.

### Ligand binding to preferential enzyme residues

The residues around 4Å of the ligand subgroups were noted for each complex. K11777-docked cryptopain-1 was taken as a reference, as K11777 had been shown experimentally (on bench) to bind Cryptopian-1. The protease subsite residues were thus primarily derived from this complex. **Figure 2** show the chosen conformation of K11777 docked into cryptopain-1 with the derived subsites colored differently. For the other best-scored complexes, the additional contact residues that showed up were assigned subsites according to their vicinity/placement to the already derived subsite residues in the three dimensional structure of cryptopain-1. **Figure 3** shows all the residues that were contacted by ligand subgroups across the enzymatic cleft, in one or more of the complexes. The panels A, B, C and D of **Figure 4** show the selected conformations of the other vinyl sulfones in the cryptopain-1, amidst the subsites derived from the reference complex.

**Figure 3:**
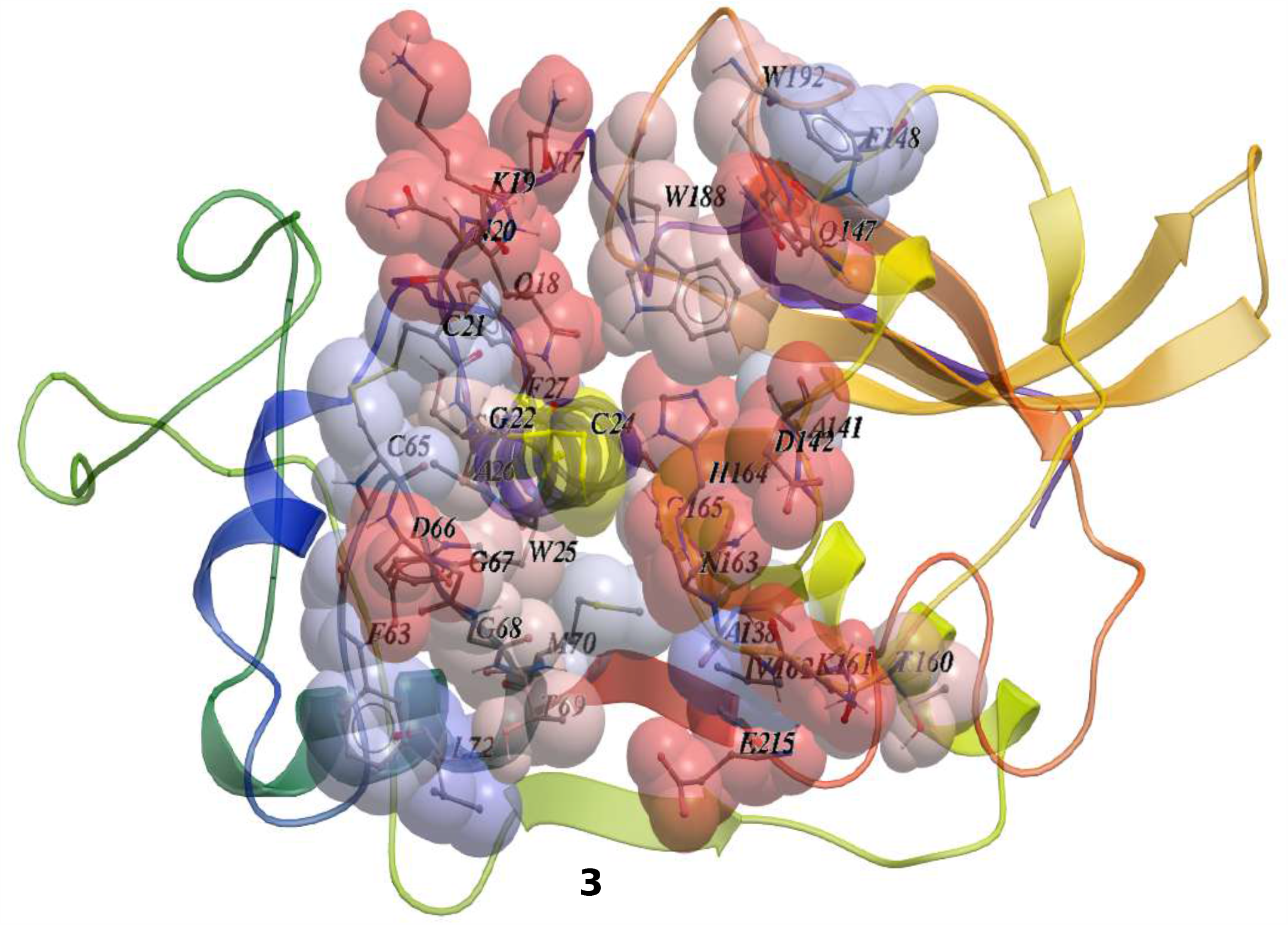
All the residues that are contacted by one or more ligands in the docked complexes of K11777 and the best-scored (score <= −29.0) vinyl sulfones are labeled and shown in spacefill representation (colored as per hydrophobicity) in the three dimensional structure (homology model) of cryptopain-1. The enzymatic triad residue C24 − the site of covalent attachment - is in yellow.

**Figure 4:**
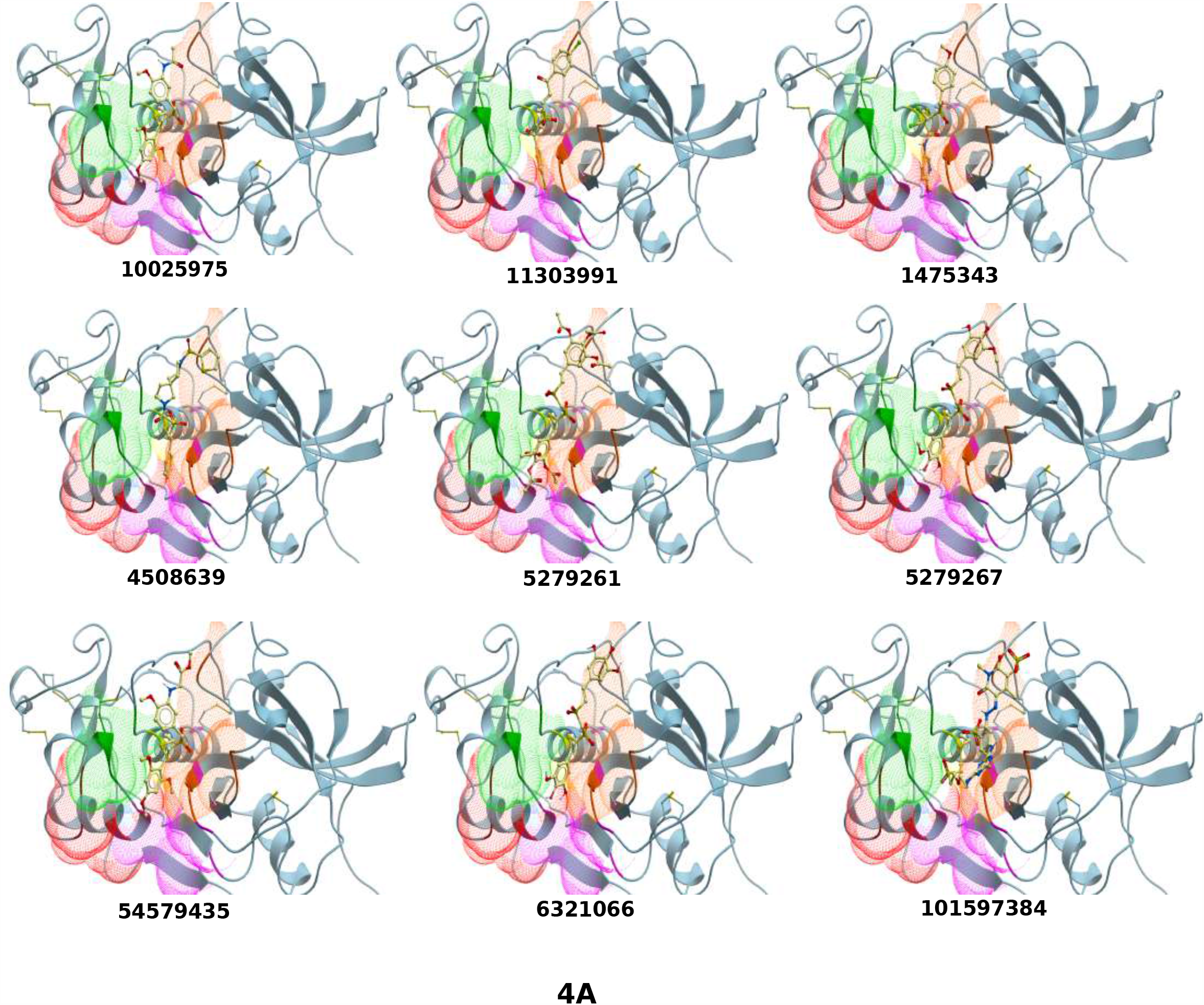
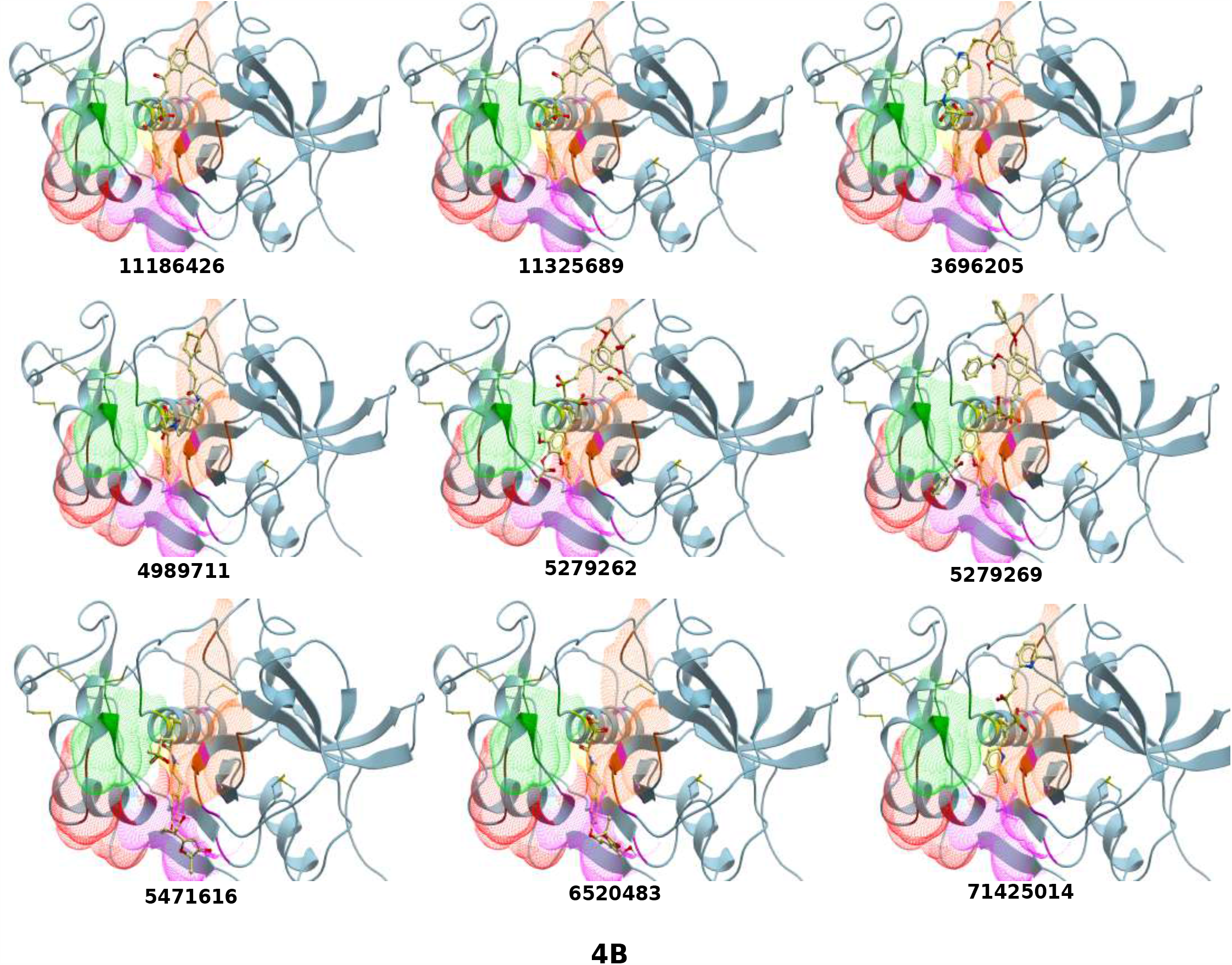
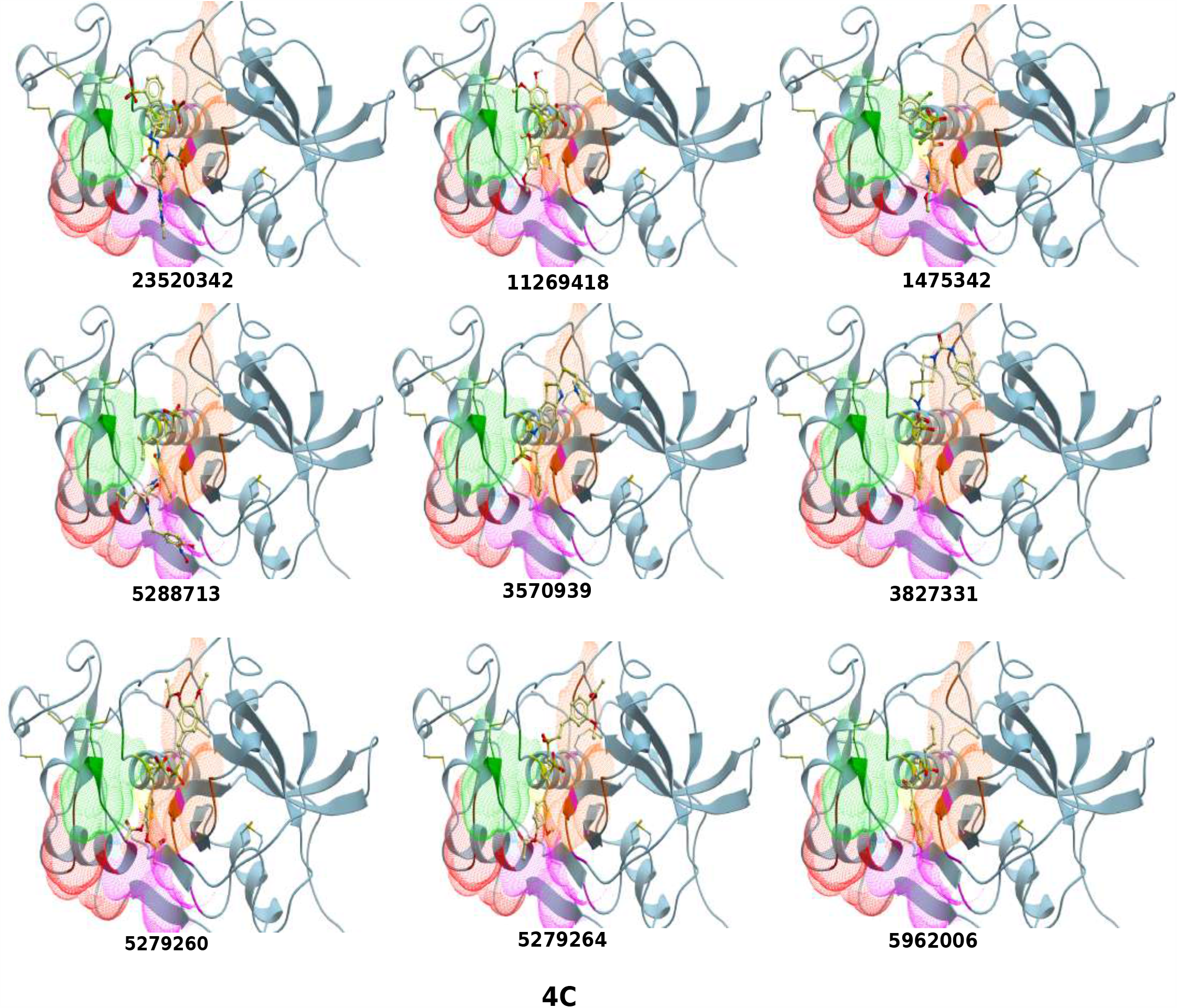
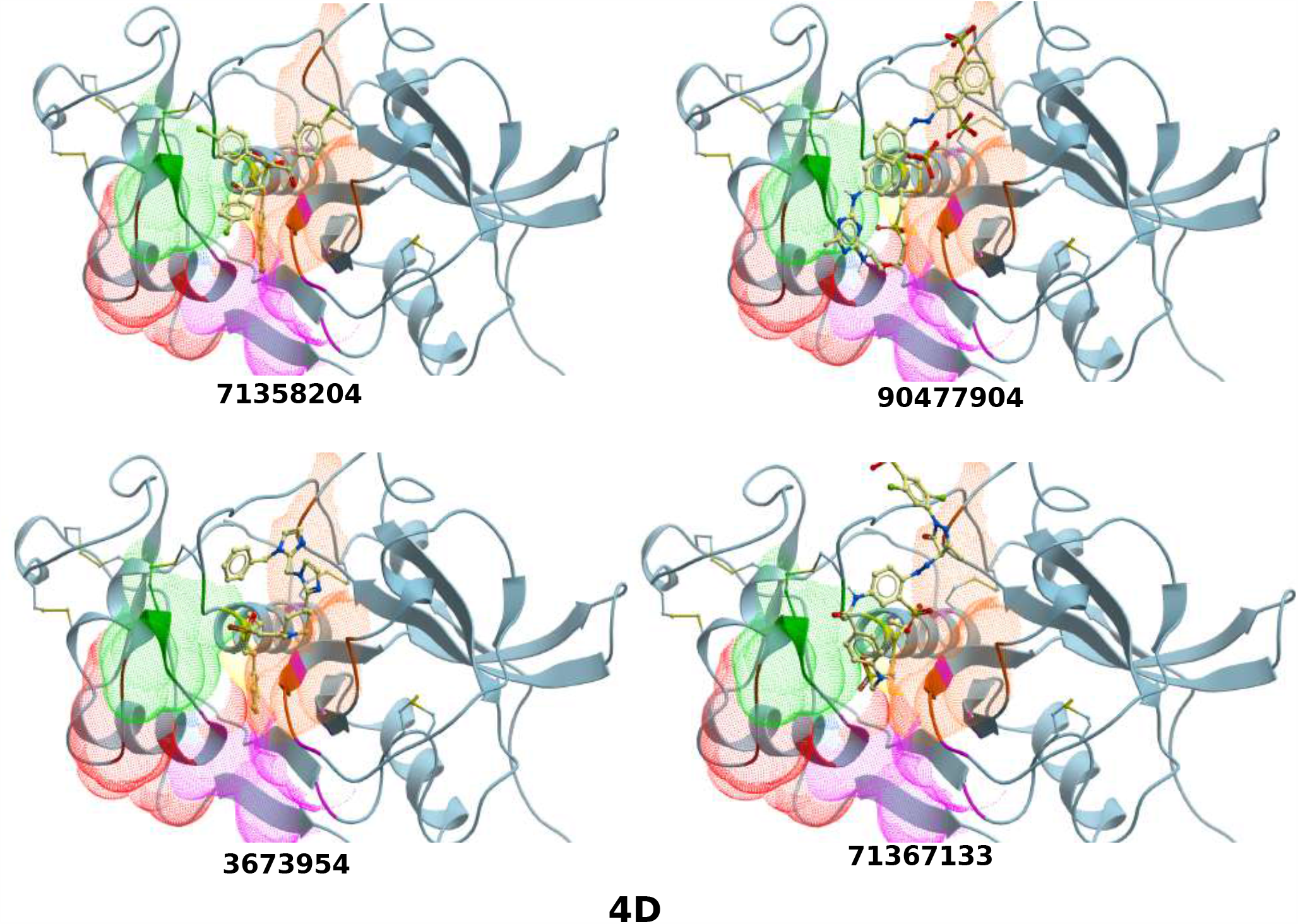
Panels A, B, C, D show the orientation and placement of the best-scored (score <= −29.0) compounds docked into the cryptopain-1 theoretical structure. The ligands are shown with respect to the enzyme subsites that have been derived from the K11777-cryptopain-1 reference complex.

The ligand subgroup-contacting residues in each complex had been mutated to Alanine; one at a time, to figure the favorable interactions based on the ddGbind values. The interactions that showed ddGbind values worse than −1 (less than −1) were not taken into account. The residues that corresponded with the rest of the ddGbind values (greater than −1) were considered to be contributing to favorable interactions with the ligand. **Supplementary Table 1** lists the ddGbind interactions in terms of residue versus ligand (represented by PubChem IDs). The columns have all the residues that had been favorably contacted in one or many of the complexes, and the rows hold the compounds whose subgroups had shown favorable interactions with the corresponding column residues. **Table 1** lists the scores, contact residues, H-bonding residues and the favorably interacting subsite residues (derived from **Supplementary Table 1**) in the complexes. The tables feature also the additional subsite residues that showed up in the other best-scored complexes, which included ligands that, unlike K11777, were not typical peptidyl vinyl sulfones.

**Table 1:**
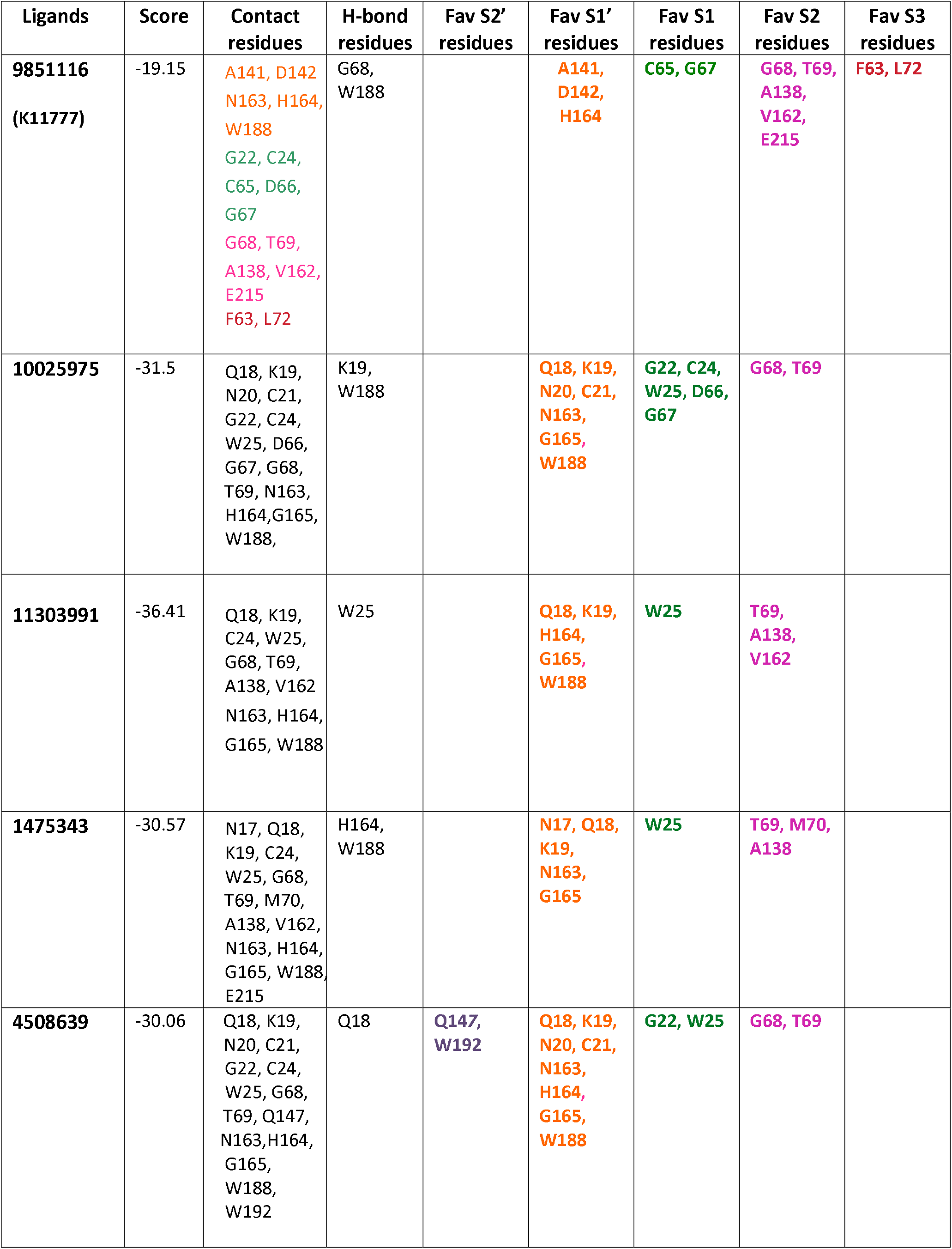
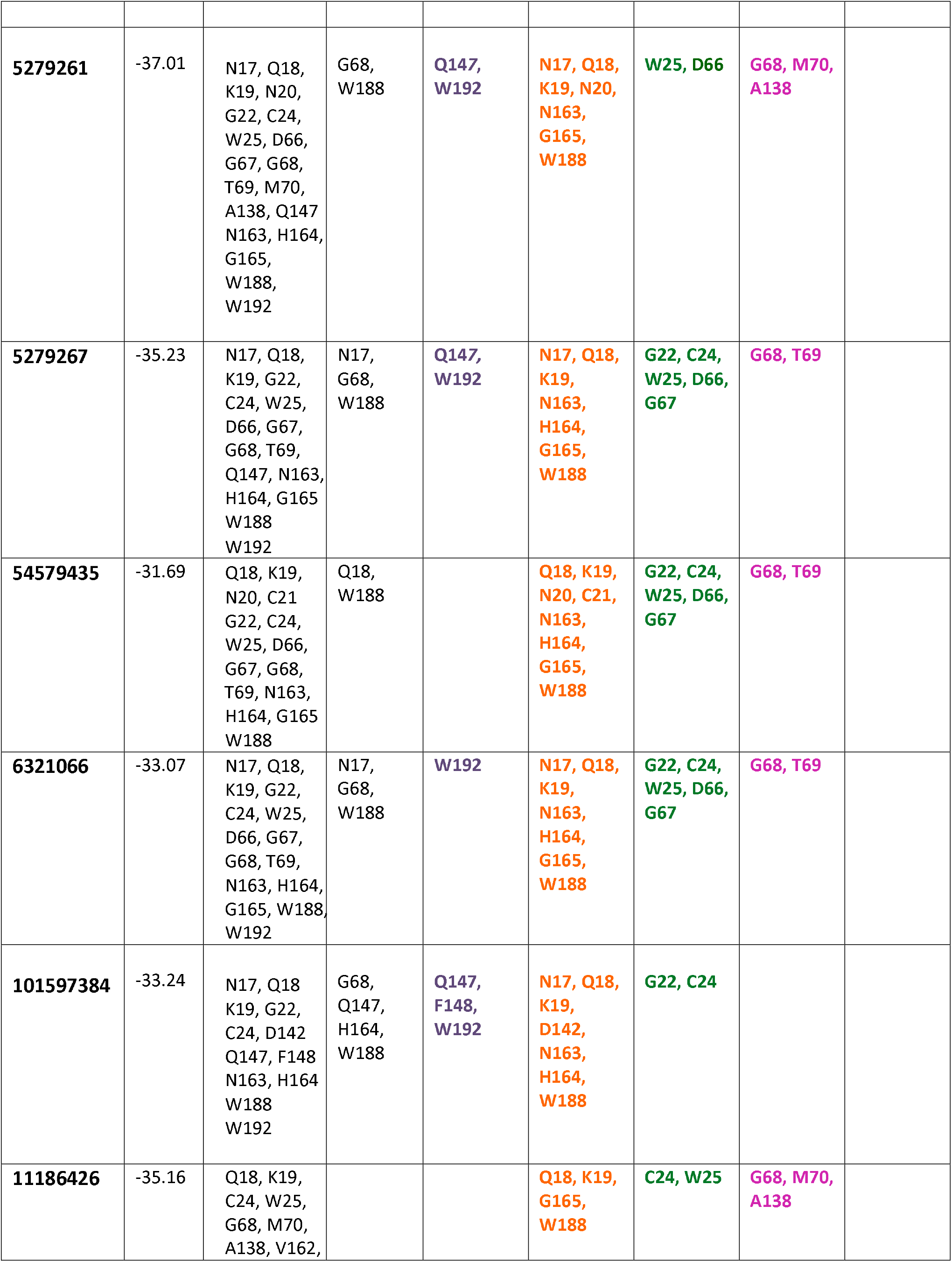
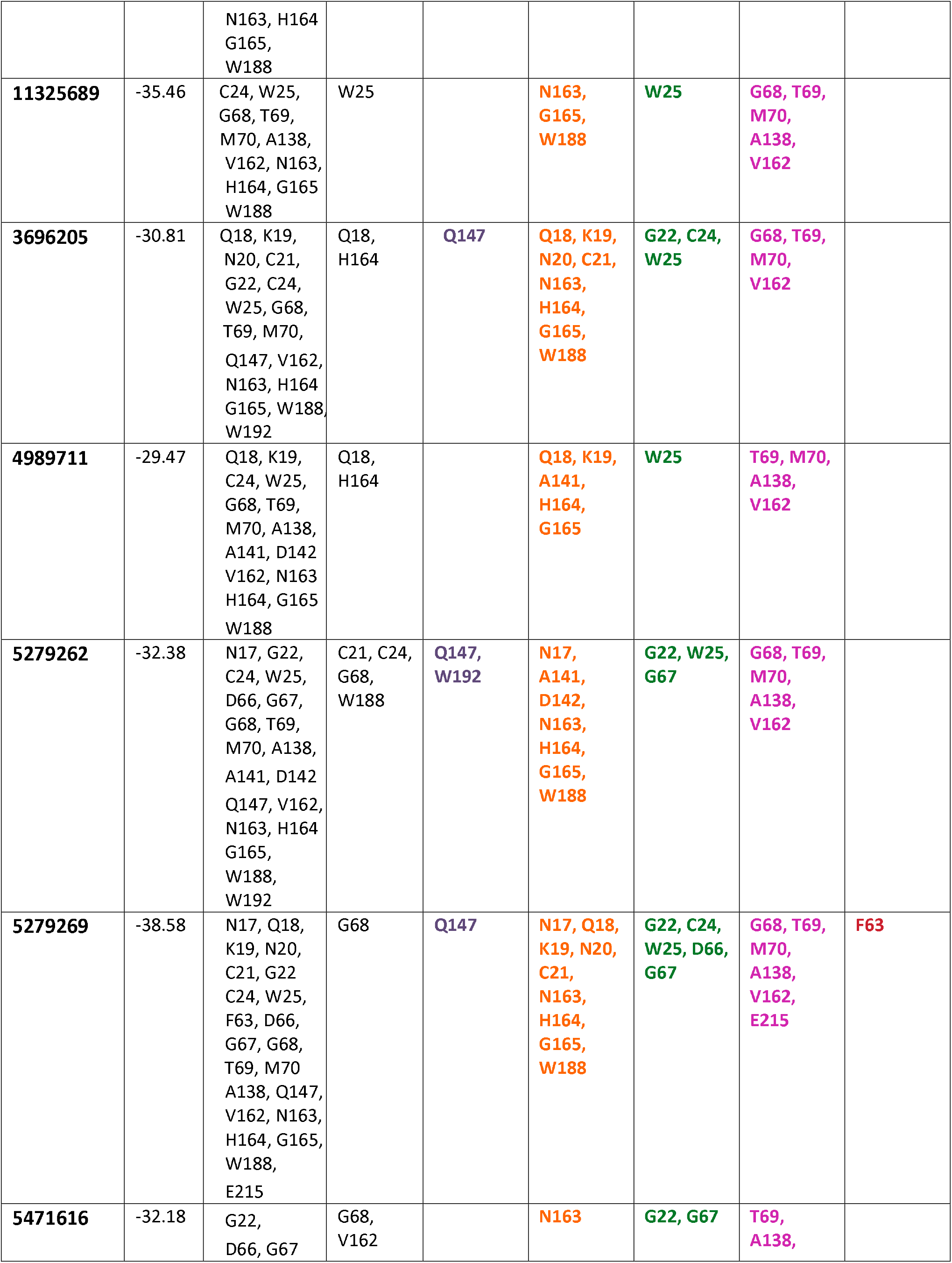
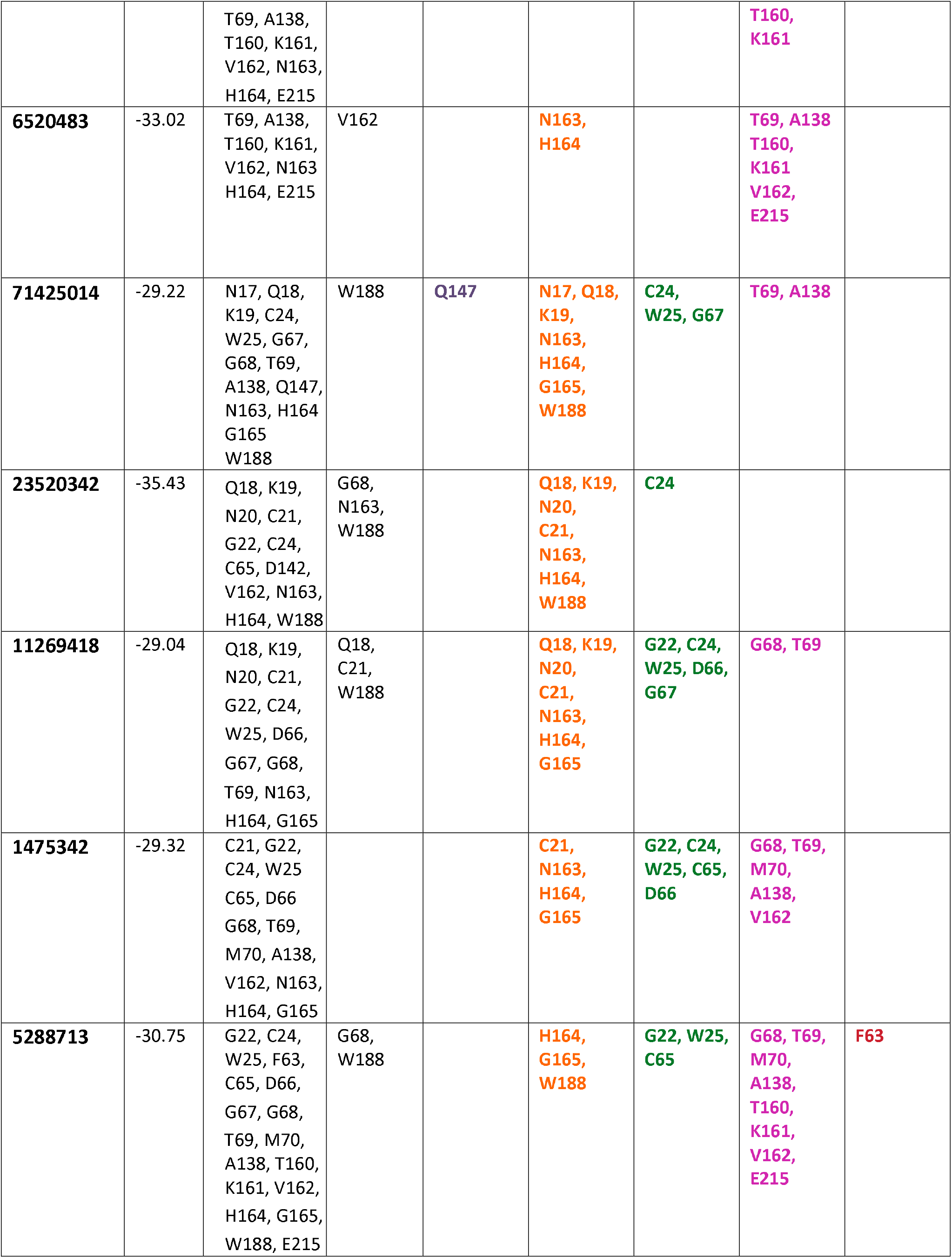
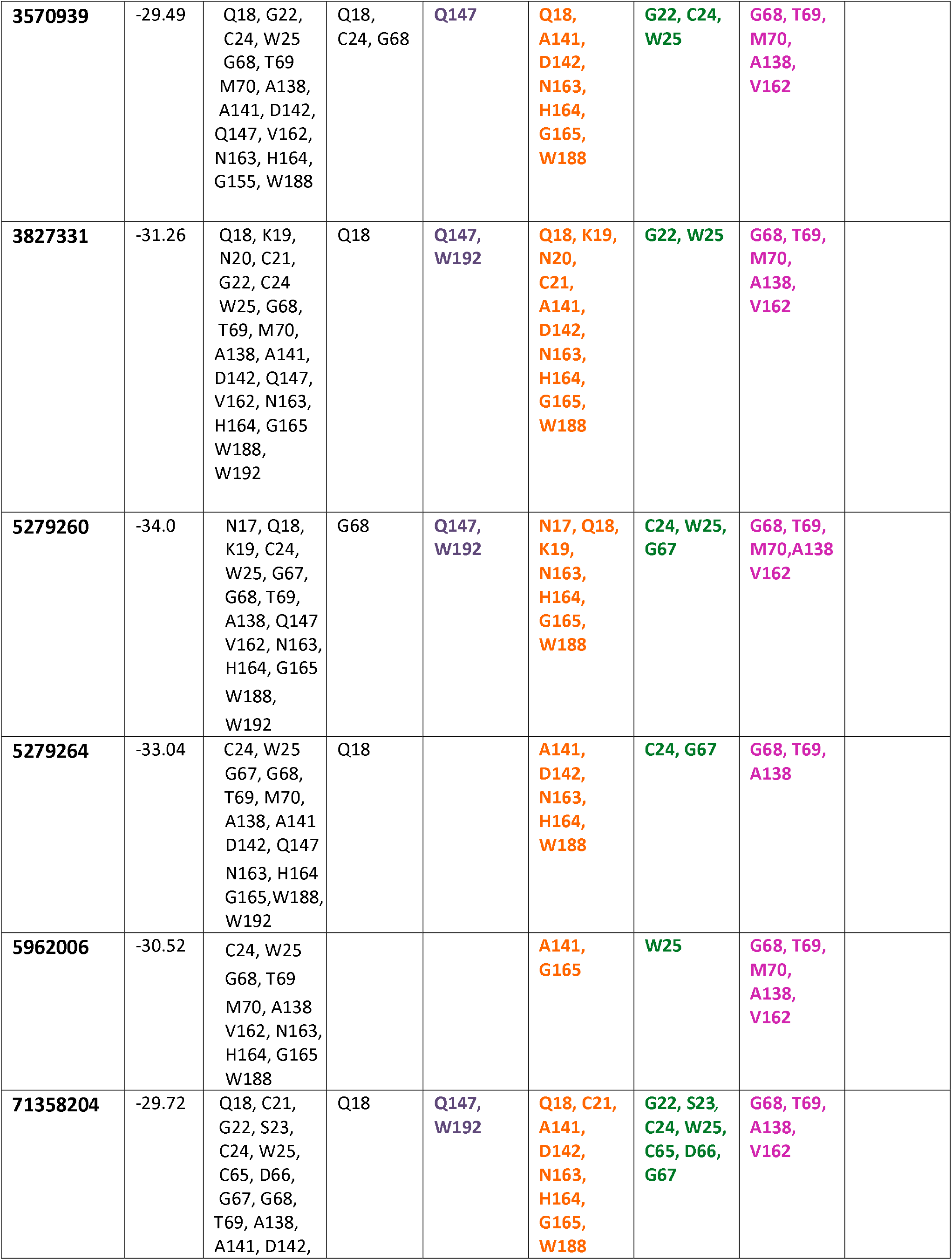
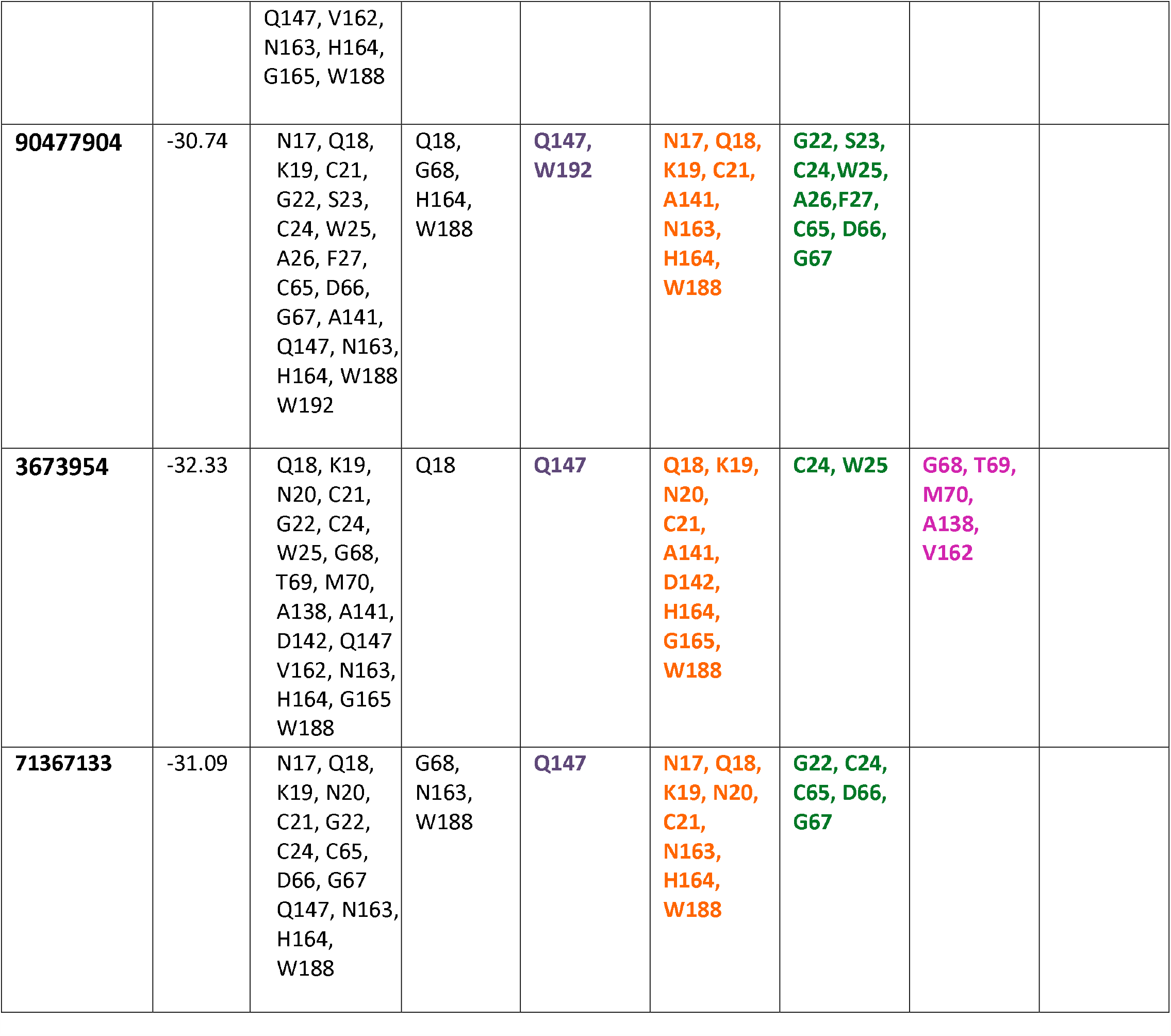
The contact residues around K11777 in cryptopain-1 are color-coded as per subsites. The residues around the P1’ sidegroup of K11777 (S1’ subsite) are in orange. The S1 site is in green, S2 in pink and S3 in red. The residues that made favorable contacts with K11777 are shown in bold in the subsequent columns. The residues around the ligand subgroups of the best-scored vinyl sulfones compounds (PubChem IDs in ligands column) are listed. The favorable interactions (including additional contact residues, which does not appear for K11777) are shown in bold and colored as per subsites. The additional S2’ subsite is shown in mauve. The scores and the H-bonding residues for the individual complexes are also listed.

Thirteen of the favorably interacting cryptopain-1 residues emerged to be heavily contacted by ligand subgroups in the complexes (see **Supplementary Table 1**). The number of times each of the residues was shown to make favorable interactions ranged from 1 to 26. With a threshold of 16, Q18, K19, G22, C24, W25, G68, T69, A138, V162, N163, H164, G165, and W188 turned out to be the most frequently contacted of the favorably interacting residues. The derived residues were then subjected to ddGbind recalculations (barring A138). The results from the calculations were studied with respect to the orientation and positioning of the ligand subgroups near the mentioned residues in the complexes. The ddGbind values for the interaction of the frequently contacted residues with the ligands are listed in **Table 2**. The purpose was to deduce the contributing factors for binding and to shed light on the enzymatic-pocket preference for accommodating certain ligand groups, which could be ultimately useful for designing a potent vinyl sulfone inhibitor (better than K11777) to target cryptopain-1.

**Table 2:** The ddGbind values for the interaction of K11777 and the best-scored ligands with the important residues of cryptopain-1 are tabulated. The residues that had showed high number of favorable interactions (Supplementary Table 1) were taken into consideration for the second round of calculations to chart this table. The values for the most favorable interactions are shown in purple, moderately favorable interactions in brown, slightly unfavorable in aquamarine and unfavorable in blue. The scale for demarcation varies for each residue, depending on the range and type of its interactions.

|  | Q18 | K19 | G22 | C24 | W25 | G68 | T69 | V162 | N163 | H164 | G165 | W188 |
| --- | --- | --- | --- | --- | --- | --- | --- | --- | --- | --- | --- | --- |
| K11777<br>(9851116) | -0.69 | 0.06 | -1.02 | -4.75 | -0.21 | -0.01 | -0.04 | -0.24 | -0.12 | -0.84 | 0.08 | -14.27 |
| 10025975 | -0.11 | 0.05 | -0.02 | -4.19 | -0.05 | 0 | -0.12 | -0.22 | -0.15 | -1.64 | 0.05 | -0.38 |
| 11303991 | -0.06 | 0.05 | -0.08 | -38.16 | 0.01 | 0 | -0.1 | 0.13 | -0.56 | -0.36 | -0.06 | -0.15 |
| 1475343 | 2.3 | 10.87 | 19.15 | -61.75 | 13.68 | -1.96 | 9.65 | 9 | -0.95 | 14.48 | 11.6 | 12.11 |
| 4508639 | 78.92 | 0.05 | -0.02 | -0.2 | -0.02 | 0 | -0.09 | -0.21 | -0.01 | -0.01 | 0.06 | 94.15 |
| 5279261 | -0.27 | -23.96 | -1.07 | -14.28 | 0.49 | 0.01 | -0.08 | -1.37 | 0.92 | -1.17 | -0.29 | 0.92 |
| 5279267 | -0.01 | 0.05 | -0.02 | 1.12 | 0.02 | 0 | -0.06 | -0.21 | 0.12 | -0.09 | 0.01 | -0.27 |
| 54579435 | -0.08 | 0.05 | -0.02 | 0.19 | -0.02 | 0 | -0.12 | -0.22 | -0.01 | -0.62 | 0.05 | -0.26 |
| 6321066 | -0.08 | 0.05 | -0.02 | 0.38 | 0.04 | 0 | -0.09 | -0.21 | -0.01 | -0.11 | 0.01 | -0.26 |
| 101597384 | -0.41 | -0.3 | -0.04 | 0.13 | -0.05 | 0 | -0.09 | -1.07 | -0.01 | -0.14 | 0.01 | -0.1 |
| 11186426 | 6.07 | 4.06 | 5.07 | 7.65 | 6.21 | 13.3 | 6.12 | 4.24 | 1.41 | 8.12 | 2.37 | 3.15 |
| 11325689 | -0.08 | 0.17 | -0.96 | -26.39 | 0.77 | 0.01 | -0.23 | 0.56 | 0.83 | -5.35 | -0.2 | 0.6 |
| 3696205 | -0.06 | 0.06 | -0.05 | 2.9 | -0.06 | 0 | -0.1 | -0.21 | -0.01 | -0.06 | 0.01 | -0.1 |
| 4989711 | 9.33 | -124.55 | -120.55 | -121.95 | -13.62 | 40.39 | 8.65 | -28.45 | -26.17 | 14.06 | 64.76 | 37.76 |
| 5279262 | -0.06 | 0.07 | -0.13 | 47.9 | 0.25 | 0 | -0.07 | -0.21 | -0.01 | -0.05 | -0.01 | -0.2 |
| 5279269 | -0.08 | 0.05 | 20.02 | 0 | 0.07 | 0 | -0.13 | -0.21 | -0.01 | -0.08 | 0.01 | -0.1 |
| 5471616 | 5.13 | 4.65 | 5.19 | 5.54 | -1.93 | 2.95 | 5.84 | 4.95 | 1.51 | 5.13 | 4.34 | 2.05 |
| 6520483 | -0.08 | 0.05 | -0.04 | 1.27 | 0.01 | 0 | -0.11 | -0.18 | 0.14 | -0.07 | 0.01 | -0.1 |
| 71425014 | -28.4 | -7.13 | -28.42 | -35.48 | 18.81 | -28.39 | -2.23 | -28.68 | 31.35 | -6.79 | 24.72 | -15.86 |
| 23520342 | 30.29 | 29.22 | 52.49 | 67.44 | 43.47 | 24.98 | 24.35 | 57.59 | 36.59 | 1.54 | 35.81 | 3.1 |
| 11269418 | -0.08 | 0.05 | -0.02 | 0.52 | -0.02 | 0 | -0.12 | -0.22 | -0.05 | -0.07 | 0.05 | -0.1 |
| 1475342 | -0.36 | 0.05 | -0.03 | 19.88 | 0.11 | 0 | -0.1 | -0.26 | -0.47 | -0.11 | -0.01 | -0.13 |
| 5288713 | 1.12 | 0.15 | -0.4 | -38.63 | -1.21 | -0.21 | -0.16 | -0.13 | -0.03 | 0.25 | 0.08 | 1.13 |
| 3570939 | -2.24 | 0.05 | -0.03 | 0.11 | -0.05 | 0 | -0.12 | -0.2 | -0.01 | -0.14 | 0.3 | -0.1 |
| 3827331 | -0.21 | 0.18 | 0.13 | -1.65 | -0.05 | 0 | -0.09 | -0.21 | 1.07 | 1.61 | -0.08 | -0.1 |
| 5279260 | -0.08 | 0.05 | -0.02 | 0.78 | 0.07 | 0 | -0.11 | -0.2 | -0.02 | -0.1 | 0.01 | -0.1 |
| 5279264 | -0.08 | 0.05 | -0.02 | 93.87 | -0.02 | 0 | -0.07 | -0.21 | -0.01 | -0.07 | 0.05 | -0.1 |
| 5962006 | 10.18 | 12.74 | -10.64 | -12.15 | 6.82 | 4.4 | 7.33 | -0.8 | -2.89 | -2.83 | 11.11 | 0.6 |
| 71358204 | -0.08 | 0.05 | -0.02 | 27.47 | 0.07 | 0 | -0.05 | -0.21 | -0.01 | -0.07 | 0.01 | -0.1 |
| 90477904 | -0.89 | -0.72 | -0.79 | 20.44 | -2.89 | 5.42 | 0.04 | -0.99 | 0.05 | -2.09 | -0.76 | 1.17 |
| 3673954 | -0.08 | 0.06 | -0.02 | 101.77 | -0.09 | 0 | -0.09 | -0.21 | -1.94 | -0.07 | -2.41 | -0.1 |
| 71367133 | 8.18 | -3.76 | -4.19 | 71.7 | 2.88 | 14.46 | -4.82 | 7.21 | 2.11 | 11.2 | 13.05 | -1.88 |

### Interactions: enzyme subsite residues - ligand subgroups

Unlike K11777 which occupied the central part of the pocket and was spread equally amongst all the subsites (**Figure 2**), the best-scored vinyl sulfones more often occupied the upper part of the cleft and tended to position themselves on the right, making contacts mostly with S1’ and S1. Ligands that lacked P1’, P2’ etc., were sometimes exceptions and got placed at the lower end of the cleft, heavily contacting S2.

The positioning of the ligand-contacting residues in the three dimensional structure of the enzyme can be seen in **Figure 3**, and the other vinyl sulfone ligands’ placement therein is visible in **Figure 4**. The accommodation of various ligand subgroups of the best-scored vinyl sulfones across the enzymatic cleft is described as follows.

### S2’ enzyme subsite

The S2’ subsite residues F148 and W192, in the uppermost part of the pocket, were not amongst the frequently contacted, and hence they were excluded from detailed analysis.

### S1’ enzyme subsite

The derived S1’ residues N163, H164 and W188 were frequently contacted by the other vinyl sulfones, along with an additional G165 (placed between N163 and H164). Q18 and K19 also featured as additional contacts, which though positioned on the opposite side in the structure, made interactions with P1’ of the ligands. Thus the residues were categorized as part of S1’.

The upper part of the heavily occupied enzymatic pocket region is constituted by S1’ residues: W188 on the right, and Q18, K19 on the left.

**W188**, which made most of the hydrophobic interactions, on the right side of the pocket, with the ligand ring systems showed highly positive ddGbind values for thiophen group in particular. The residue seemed to prefer pi stacking with ligand ring systems as it showed favorable ddGbind values for in-plane ring interactions. The ligands with ethenyl group as well as the ones that did not place any subgroups near the residue showed moderately favorable interactions. The ligands whose rings were out of plane with the residue’s six-membered ring, and the ones that had groups like bromopyridine near the residue, showed unfavorable interactions.

For **Q18** that is situated at the back of the cleft wall, the compounds’ covalent moiety with their sulfonyl group and/or benzyl/phenyl ring(s), when placed near the lower end of the residue, resulted in favorable interactions. Large halide containing subgroups such as bromopyridine resulted in unfavorable interaction.

**K19**, positioned at the front of the cleft, showed favorable interactions with reasonably distanced polar substituents. Interactions were favorable even when no substituent was close to the residue. Understandably, unfavorable interactions were observed when the non-polar moiety of the residue’s sidechain was near polar ligand atoms, and interactions of non-polar ethenyl group of the ligand with polar end of the residue also led to highly negative ddGbind values.

The mid-region of the highly occupied cleft is constituted by N163, H164 and G165 (S1’ residues) on the right. These frequently contacted residues were actually within the contact range of both P1’ and P2 of K11777. However, the proximity of the ligand’s P1’ to the sidechains of N163 and H164 in the reference complex led to the residues’ allocation to S1’ – which therefore extends into the middle of the cleft.

**N163** showed favorable interaction with halide-containing substituents including bromopyridine that otherwise had unfavorable interactions with the other residues. The ligands that had their benzyl/phenyl rings at a comfortable distance from the residue showed favorable interactions. Closely spaced ligand ring systems led to clashes.

**H164**, which is situated at the back (compared to N163) of the enzyme’s mid-pocket, preferred favorable interactions with the ligands’ sulfonyl or backbone. The residue, if not always, showed favorable interactions even when no ligand group was placed near it. Favorable ring interactions were observed when the ligands’ ring systems were mostly tilted towards W188. Unfavorable ddGbind values were observed for inflexible ethenyl groups in ligands.

**G165**, which is buried in the mid-pocket, made interactions primarily with the covalent-bond forming moieties of the ligands. The residue showed favorable interactions with reasonably distanced ring systems. Interactions were unfavorable for closely spaced rings and inflexible groups such as ethenyl.

Overall, the arrangement of the mentioned residues suggest that substituted benzene/napthalene ring systems could be accommodated in the upper region of the subsite, where the ligand rings can engage in hydrophobic interaction with W188, and the polar substituents on those rings could interact with Q18 and K19 to the left of the pocket. However, large (polar) halide-substituted rings such as bromopyridine could lead to clashes. The S1’ in the mid-pocket shows a preference for reasonably distanced ring systems and halide-substituted ligand subgroups. The subsite is not likely to tolerate inflexible groups such as diazospiro, ethenyls etc.

### S1 enzyme subsite

The frequently contacted (derived) S1 residues G22 and C24 were positioned on the left side of the mid-pocket. W25, that emerged as an additional frequent contact was placed close-by to G22 and C24 on the left, and formed part of S1.

**G22** was observed to like interactions with double ring systems such as substituted napthalene or two separate benzyl/phenyl rings placed near the residue. It also showed favorable interactions with groups like sulfonyl and/or polar backbone atoms. Ring as well as polar interactions showed the most favorable ddGbind values. The interactions became unfavorable when no ligand group was in the vicinity of the residue. Bromopyridine showed unfavorable interactions with this residue too.

**C24**, the enzymatic triad residue that formed the covalent bond with the vinyl sulfones, preferred the ligands to be placed away from it and towards the front of the cleft. The favorably interacting compounds were positioned to the right and at the bottom of the residue. The compounds that were tilted towards the inside of the cleft showed moderately unfavorable interactions, and so did the ones that did not place any ring system near the residue. Unfavorable interactions for the residue were observed with the close proximity of ligands’ polar substituents or backbone. Again, bromopyridine made unfavorable interactions with this residue as well. Unlike other residues, C24 had far less borderline interactions and the individual ddGbind values mostly ranged on either side of favorable and unfavorable.

**W25** made favorable interactions with the ring systems of the ligands that were placed away, and towards the right side of the pocket. The interactions were better with more number of rings. The highest ddGbind value was obtained for the compound that had four ring systems. However close interactions either with the ligand backbone or side chain resulted in unfavorable interactions. Inflexible groups such as diazospiro, even if placed away from the residue, amounted to negative ddGbind values.

Taken together, inflexible groups such as diazospiro, ethenyl etc. would not be tolerated by S1. The subsite can accommodate multiple ring systems. The mid-pocket would have a preference towards polar backbone of ligands that are positioned towards the front. The catalytic C24 of S1 too dictates the compounds to be placed not too deep inside the cleft. Large halide containing subgroups such as bromopyridine will not be favored in the subsite. The site shows a propensity towards closely packed ring interactions.

### S2 enzyme subsite

The lowest part of the heavily occupied pocket is comprised by the frequently contacted (derived) S2 subsite residues: G68, T69, A138 and V162. The S2 residues are distributed on both sides of the cleft. G68, T69 are on the left, and A138, V162 are on the right.

**G68**, placed above T69, engaged mostly in H-bond interactions with backbone of the ligands, rather than favorably accommodating their side chains. The residue showed favorable ddGbind values for slightly spaced away ring systems of ligands. The most unfavorable interactions were shown for the compound containing bromopyridine.

For **T69**, the highest positive ddGbind value was observed for a halide-substituted ligand subgroup (fluro-triazinyl group) with its polar ring and polar backbone near the residue. T69 preferred reasonably distanced ring interactions (polar and non-polar). However, with no ligand group placed near the residue, the interactions were unfavorable. Also, with large subgroups like bromopyridine again, the interactions were unfavorable.

**A138** had to be excluded from the mutational analysis as ddGbind value for Ala to Ala mutation is zero, and could not have provided any useful clue towards the type of interactions.

**V162**, despite being mostly hydrophobic, showed favorable interactions with comfortably distanced polar subgroups of ligands including the fluro-triazinyl group-containing compound that showed the best ddGbind value. Such polar groups were presumably stabilized by long-ranged electrostatic effect of other S2 residues (see tables).

Summing up, S2 can certainly accommodate polar subgroups/backbone of ligands. The subsite however, like the other subsites, does not like to accommodate large polar subgroups like bromopyridine.

### Orientation and placement of ligands across the enzymatic cleft

The best-scored vinyl sulfones tended to occupy the S2’, S1’, S1 and S2 subsites. Unlike K11777, the other compounds showed optimal interactions mostly with the prime site residues of the enzyme. The S1’ residues made half of the frequently contacted favorable interactions with the ligands. The rest half of such interactions were accounted by S1 and S2 members.

With respect to the entire enzymatic cleft of cryptopain-1, it can be deduced that the ligands’ placement towards the front of the cleft would be preferred to deep-seated interactions. Polar backbones of ligands (even if not peptidyl) would be desired. S1’ and S2 like to be occupied, and are prone to make favorable interactions with polar subgroups of ligands. Large halide-containing subgroups are not well tolerated presumably because of their size. Reasonably distanced ring interactions would be preferred all across the cleft. Unlike inflexible groups like substituted napthalene which could be favorably accommodated in S1, the strain arising out of the inflexibility of ethenyl and/or diazospiro groups is not likely to be tolerated, especially in the S1’ and S1 subsites, as per the computational mutational analysis.

Quite relevantly, the compound 23520342 that showed the maximum number of favorable interactions with the frequently contacted residues, (see **Table 2**) had all the preferred attributes and lacked the undesirable ones. The ligand-bound protease showed a very good score of −35.43.

Some other compounds that showed slightly better scores than 23520342 were 11303991 (score: −36.41), 5279261 (score: −37.01), and 5279269 (score: −38.58).

11303991 and 5279261 were placed deep inside the cleft that led to clashes with the covalent bond forming C24. The ligands’ polar backbones, in addition to the occupation of the enzymatic S1’ site with polar subgroups, somewhat mitigated the unfavorable interactions in totality. The compounds also had the undesirable ethenyl near S1’, which contributed to unfavorable interactions with K19 in case of 5279261 (where the ethenyl was placed much closer to the residue). However, the overall scoring algorithm did not penalize ethenyl’s presence as much as the individual ddGbind calculations did.

5279269, which showed the best score, too had an ethenyl group (albeit not close to K19). This compound however was placed towards the front of the cleft, thereby avoiding unfavorable interactions with C24. Also, the ligand had ring systems in abundance (six) for favorable interactions. Rings comprised its (polar) backbone as well as subgroups. The ligand desirably occupied the S1’ and S2 subsites, though not with much polar subgroups.

## CONCLUSION

The efficacy of the thirty-one best-scored compounds as drug candidates within physiological limits remains to be tested on bench. The information, which has been garnered through this study on the substrate/ligand-binding cleft of the enzyme and its interaction with the chemical groups of the docked compounds, could ultimately guide the design of potent vinyl sulfone inhibitors.

23520342 and 5279269 that shared most of the preferred ligand-subgroup attributes can serve as model compounds, based on which effective inhibitors against cryptopain-1 could be designed. **Figure 5** provides the chemical structures of the reference (K11777) and the model compounds. Unlike the other two mentioned compounds (11303991 and 5279261), the subgroups of the model ligands extended into S2 – typically the key specificity determinant in cathepsin L-like cysteine proteases such as cryptopain-1. 23520342 placed a polar subgroup at S2 in contrast to the hydrophobic subgroup put by 5279269. Polar ligand subgroups (as in 23520342) at the enzyme’s S2 are likely to be stabilized via polar/electrostatic interactions by residues like T69, M70, T160, K161 and E215. Hydrophobic subgroups too (as in 5279269) could be accommodated by the virtue of S2 residues like A138 and V162.

**Figure 5:**
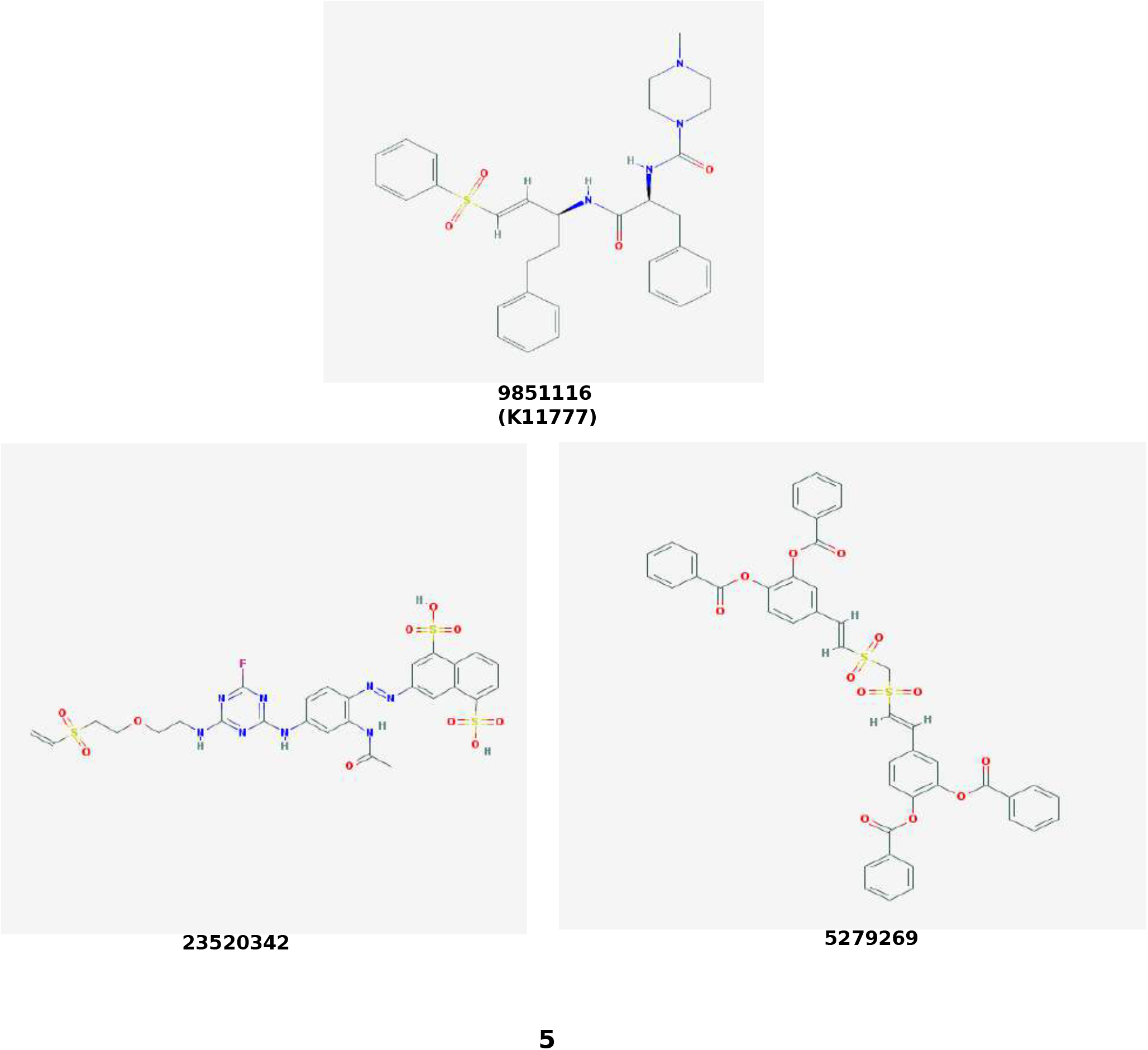
The chemical structures (along with the PubChem identifiers) of the reference ligand K11777 or K-777, and the two model compounds - which showed optimum interactions with the enzymatic cleft of cryptopain-1 and thereby could aid the design of effective inhibitors to target the protease.

Thus, the study attempted to identify purchasable vinyl sulfone compounds that can possibly inhibit cryptopain-1, as well as it provided crucial information pertaining to receptor-ligand interactions to help future design of other vinyl sulfones, which could prove to be effective in curbing cryptosporidiosis.

## Acknowledgement

The author would like to thank Prof. Ruben Abagyan of University of California San Diego, for providing computational resources.

