## supplementary figures for "Identification and design of vinyl sulfone inhibitors against Cryptopain-1 – a cysteine protease from cryptosporidiosis-causing *Cryptosporidium parvum*"

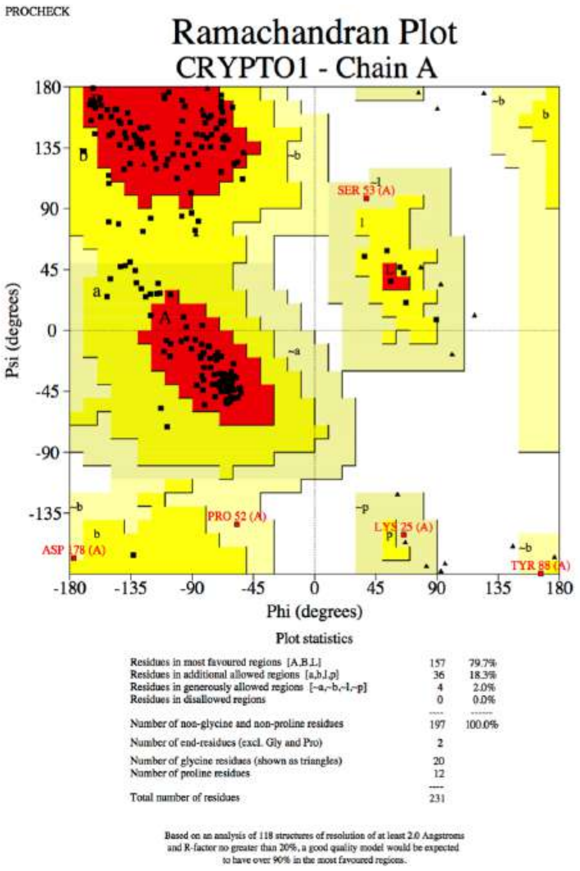


**1A**

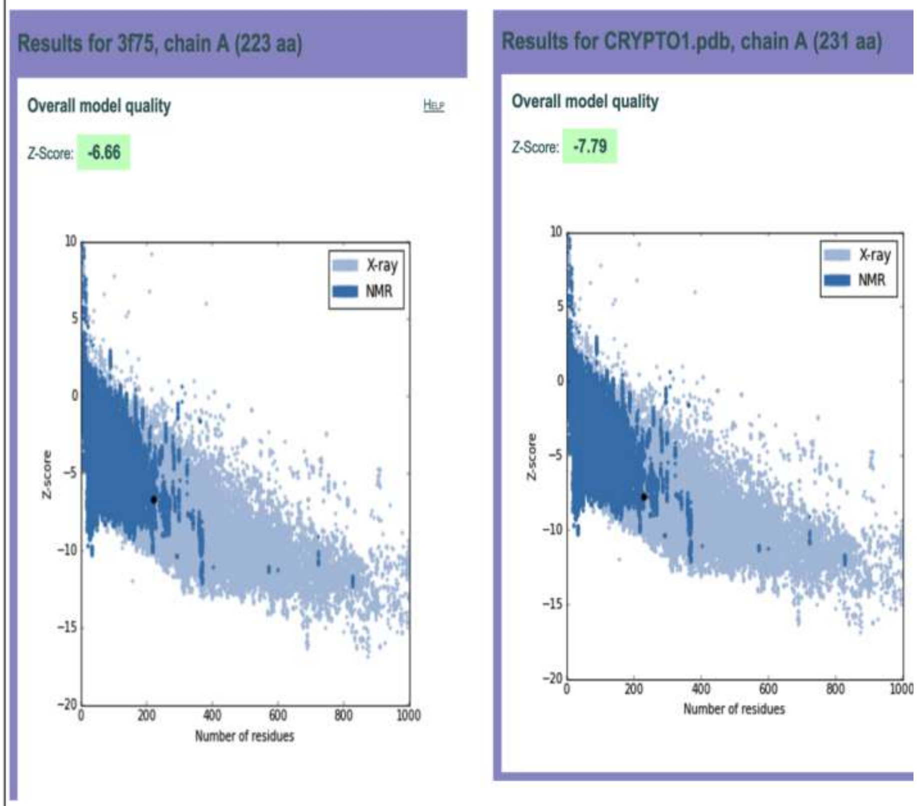


**1B**

**Figure 1A**: The ramachandran plot for the homology model of cryptopain1; **1B:** The PROSA scores for the structural quality of the template (PDB ID: 3f75) versus the cryptopain-1 homology model

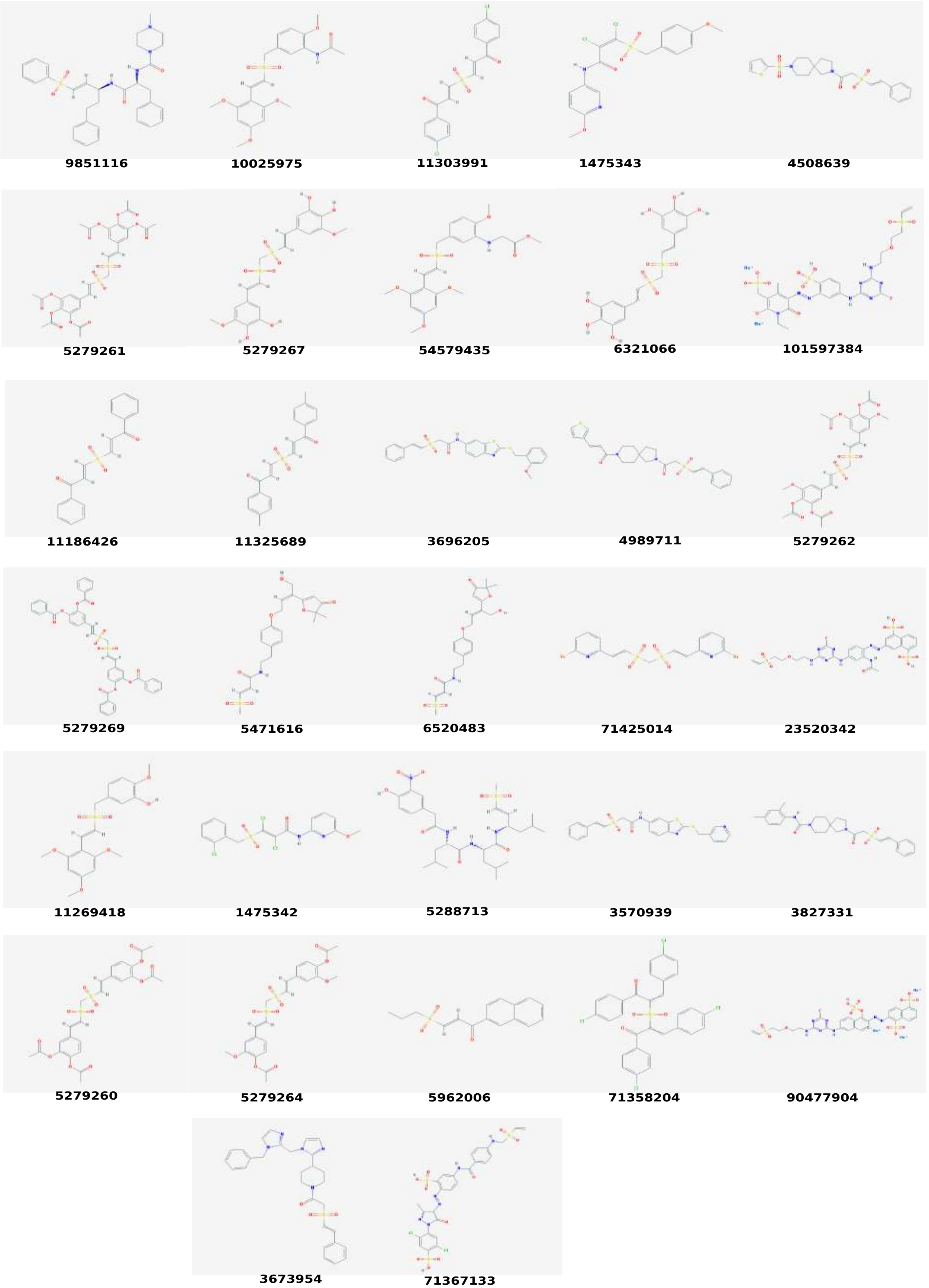


**Figure 2:** The ligand chemical structures along with the PubChem IDs of the best-scored vinyl sulfones. The IDs are from the most recent records of PubChem (as per May 26, 2018)
